# Comparison of HLA ligand elution data and binding predictions reveals varying prediction performance for the multiple motifs recognized by HLA-DQ2.5

**DOI:** 10.1101/2020.07.14.202895

**Authors:** Zeynep Koşaloğlu-Yalçın, John Sidney, William Chronister, Bjoern Peters, Alessandro Sette

**Affiliations:** La Jolla Institute for Immunology, La Jolla, CA; Department of Medicine, University of California, San Diego, La Jolla, CA

## Abstract

Binding prediction tools are commonly used to identify peptides presented on MHC class II molecules. Recently, a wealth of data in the form of naturally eluted ligands has become available and discrepancies between ligand elution data and binding predictions have been reported. Quantitative metrics for such comparisons are currently lacking. In this study, we assessed how efficiently MHC class II binding predictions can identify naturally eluted peptides, and investigated instances with discrepancies between the two methods in detail. We found that, in general, MHC class II eluted ligands are predicted to bind to their reported restriction element with high affinity. But, for several studies reporting an increased number of ligands that were not predicted to bind, we found that the reported MHC restriction was ambiguous. Additional analyses determined that most of the ligands predicted to not bind are either weak binders or predicted to bind other co-expressed MHC class II molecules. For selected alleles, we addressed discrepancies between elution data and binding predictions by experimental measurements, and found that predicted and measured affinities correlate well. For DQA1*05:01/DQB1*02:01 (DQ2.5) however, binding predictions did miss several peptides that were determined experimentally to be binders. For these peptides and several known DQ2.5 binders we determined key residues for conferring DQ2.5 binding capacity, which revealed that DQ2.5 utilizes two different binding motifs, of which only one is predicted effectively. These findings have important implications for the interpretation of ligand elution data and for the improvement of MHC class II binding predictions.

## INTRODUCTION

Major histocompatibility complex (MHC) class II molecules are expressed on professional antigen presenting cells, and present antigenic peptides to CD4+ T cells, which play an important role in regulating immune responses(1). MHC class II, called Human Leucocyte Antigen (HLA) class II in humans, and CD4+ T cell responses are of major significance for autoimmunity and antitumor immunity. It is known, for example, that celiac disease is driven by CD4+ T cell responses to gluten proteins presented by certain HLA class II molecules(2). More recently, it has been shown that neoantigens presented on HLA class II elicit potent antitumor responses in patients and that CD4+ T cells are required for successful anticancer immunotherapy (3–5). Given these developments, there is an increased interest in developing efficient approaches for identifying epitopes presented by MHC class II. As experimental identification of epitopes remains time consuming and costly, and subject to several technical challenges, MHC class II binding prediction tools are now commonly used(6).

Most MHC binding prediction tools are based on machine learning algorithms trained on datasets of known peptide–MHC binding affinities. With an increasing amount of data available for training, prediction algorithms have been able to achieve high accuracy. It is however generally accepted that predicting binding to MHC class II is a more challenging problem when compared to predicting binding to MHC class I due to a number of reasons. MHC class II molecules consist of heterodimers of alpha and beta chains. These chains are encoded by genes in the HLA-DP, -DQ, or -DR region of chromosome 6 in humans and are highly polymorphic in the general population (7). Most of the polymorphism is associated with residues forming the MHC’s peptide-binding region, which accounts in large part for the high degree of variation in binding specificity observed between different alleles. The presence of two potentially polymorphic chains makes the assignment of MHC restriction harder for class II, compared to class I molecules, which consist of a variable alpha chain and the monomorphic beta-2-microglobulin molecule.

The main energy of interaction for peptide binding to MHC class I and II is conferred by a peptide core of approximately 9 residues. However, in contrast to MHC class I, the binding groove of MHC class II is open on both ends, allowing peptides of variable length to bind. The presence of amino acids flanking the peptide core is also necessary for efficient binding to MHC class II. As a result, MHC class II have a preferred minimal length of about 13 residues for binding peptides. The presence of such over-hanging residues, plus the fact that the open ends of the binding groove afford multiple different peptide frames, means that machine learning algorithms for MHC class II binding have to consider multiple possible peptide cores.

The specificity of an MHC molecule can be described as a ‘motif’. Crystallographic analyses of MHC-peptide complexes have shown that different MHC molecules have different pockets within their binding groove. When a peptide is bound, the amino acid side chains of so-called anchor residues in the peptide specifically insert into these pockets within the MHC binding groove. Both the position and specificity of these anchor residues are similar for different peptides binding the same MHC molecule and together describe the peptide-binding motif of a given MHC molecule. For MHC class I binding peptides the anchor residues are located at or near the N- and C-termini of the peptide, usually P2 and P9 in the case of 9-mer peptides. For MHC class I binding peptides anchor residues can be at various distances from the ends of the peptide and at various positions of the binding core. For DRB1*01:01 and DRB1*15:01, for example, the prototypical anchor positions are at P1, P4, P6, P7, and P9. But other molecules, such as DRB1*11:01 and DQA1*03:01/DQB1*03:01, appear to have slightly different anchor spacing, with the second anchor at P3 instead of P4, or an anchor at P5 (8). While for many MHC class I and class II molecules distinct binding motifs could be defined, there are also instances of MHC molecules that carry multiple binding motifs, such as HLA-DRB1*03:01 (9).

Recent advances in mass spectroscopy (MS)-based techniques has enabled the development of high-throughput ligand elution assays, which allow the identification of thousands of natural ligands with a single experiment (10, 11). As eluted ligands undergo the natural processing and MHC loading cascade, the resulting elution data will inherently contain valuable biological information that is not available when only peptide binding is considered (6). Given these advantages, ligand elution experiments have become quite popular, leading to a wealth of data in the form of eluted ligands from both MHC class I and class II molecules.

As eluted ligands were isolated from MHC molecules that presented them on the cell surface, their capacity to bind MHC is inherently supported. With the increasing amount of ligand elution data, discrepancies between MHC class II ligand elution and binding predictions have been reported and the efficiency of binding predictions has been put into doubt (12–15). However, quantitative metrics for such comparisons are currently lacking. In this study, we wanted to systematically investigate how well HLA class II binding predictions and ligand elution data agree with each other. To achieve that, we assembled HLA class II ligand elution data reported in literature and performed multiple comparative analyses between elution data and binding predictions: 1) We analyzed how many of the eluted ligands were predicted to bind their reported restricting MHC 2) For several MHC class II alleles a significant fraction of reported ligands were not predicted to bind and we investigated these in detail 3) We then chose four representative HLA class II alleles and specifically selected peptides for which ligand elution data and binding predictions disagreed and experimentally measured binding affinity to determine which method was correct 4) We found that the correlation between predicted and measured binding is lowest for DQA1*05:01/DQB1*02:01 (DQ2.5). We therefore investigated this allele in detail and performed experiments to the determine key residues for binding. We found that DQ2.5 can bind peptides using alternate binding modes and that currently, only one of these is being predicted accurately.

## RESULTS

### HLA class II reported eluted ligands are, in general, predicted to bind to their restriction element

We first addressed whether ligands eluted from a specific HLA would also be predicted to bind that allele. We utilized 1,000 nM as an affinity threshold to classify peptides into binders and non-binders. This threshold was chosen based on multiple previous experimental studies analyzing the binding affinity of HLA class II restricted T cell epitopes (16–18). These studies suggested that 1,000 nM is a generally applicable threshold for biologically relevant binding in the context of HLA class II. This threshold was also confirmed in a recent analysis which found that 83.3% of HLA class II restricted epitopes are predicted to bind their corresponding restricting allele with an affinity of 1,000 nM, or better (19). Of note, optimal thresholds for MHC class II are still unclear and are being discussed in the community. The 1,000 nM threshold is a rather comprehensive threshold. In this study, we wanted to assess the binding capacity of reported eluted ligands which is why a comprehensive threshold was suitable. In a setting where the rationale is to use binding predictions to select the best binders for experimental testing, another strategy would be recommended, such as selecting the top 10% of predicted binders.

To address the extent to which HLA class II eluted peptides are predicted by the 1,000 nM threshold, we used the Immune Epitope Database (IEDB) and downloaded all peptides reported as eluted from HLA class II expressing cells. We retained peptides of 15-20 amino acids in length, and eluted from HLA molecules typed to the 4-digit resolution level. We only further considered alleles for which at least 10 ligands were reported. The resulting dataset included 12,449 sequences eluted from 16 different HLA class II molecules.

Next, for each of these 16 HLA class II molecules and their associated ligands, we performed predictions using the NetMHCIIpan algorithm (20, 21). As was the case with T cell epitopes (22), significant variation in prediction performance across alleles was noted. On average, 64% +/−31% SD (**Table 1**) of eluted peptides were predicted to bind the corresponding allele at the 1,000nM level.

**Table 1.** Eluted ligands by HLA class II molecule

| **HLA allele** | **No.**<br>**ligands** | **%recovered**<br>**(1000 nM)** |
| --- | --- | --- |
| DRB3*02:02 | 10 | 100 |
| DRB1*01:01 | 594 | 95 |
| DRB1*04:04 | 35 | 89 |
| DRB5*01:01 | 131 | 88 |
| DRB1*11:01 | 50 | 86 |
| DQA1*05:05/DQB1*03:01 | 3908 | 85 |
| DQA1*01:02/DQB1*06:02 | 27 | 85 |
| DRB1*04:05 | 1206 | 80 |
| DRB1*15:01 | 330 | 74 |
| DRB1*04:01 | 463 | 63 |
| DRB1*03:01 | 79 | 52 |
| DQA1*05:01/DQB1*02:01 | 2055 | 45 |
| DQA1*01:01/DQB1*05:01 | 23 | 35 |
| DRB1*13:02 | 17 | 35 |
| DQA1*02:01/DQB1*02:02 | 3492 | 8.0 |
| DRB1*04:02 | 29 | 3.0 |

Thus, while the majority of eluted peptides could be effectively predicted by existing algorithms, the overall prediction efficiency was lower than what has been reported for known T cell epitopes. Further, the data also revealed significant inter-allele variability, and identified several alleles (**Table 1**), including HLA-DQA1*02:01/DQB1*02:02, DQA1*05:01/DQB1*02:01, DQA1*01:01/DQB1*05:01, DRB1*04:01, DRB1*03:01, DRB1*04:02, DRB1*13:02 and DRB1*15:01, for which less than 75% of reported eluted ligands were predicted to bind at the 1,000nM level.

### Complexity and limitations of eluted ligand determinations

Of the eleven DR molecules studied, four (DRB1*04:01, DRB1*03:01, DRB1*04:02, DRB1*13:02) were associated with lower prediction efficiency. We suspected that the poor-performing DR alleles may have resulted from an imprecise HLA assignment of the eluted ligands. To further explore this issue, we scrutinized papers that reported the eluted ligands associated with those DR molecules. HLA-DR loci encompass a monomorphic alpha chain (DRA), which pairs with beta chains expressed by two different loci: DRB1 and DRB3/4/5. These two loci are in strong linkage disequilibrium and are co-expressed *in vivo.* The only exceptions to this are the DR1, DR8 and DR10 haplotypes, for which there is no associated DRB3/4/5locus. Thus, in heterozygous individuals, up to four different DR molecules (two with DRB1- and two with DRB3/4/5 beta chains) may be expressed, and even in homozygous humans, two different DR molecules may be co-expressed. Importantly, all anti-DR antibodies (L243, LB3.1 and others) commonly used to purify DR molecules and isolate eluted ligands are directed against the DR alpha chain, and thereby do not distinguish between DRB1 from DRB3/4/5 molecules. Thus, unless HLA mono-allelic lines are used, both DRB1 and the DRB3/4/5 gene products will be co-purified, and the pool of eluted ligands may contain a mixture of two different HLA class II molecules.

All studies pertaining to those DR alleles associated with “lower efficiency of binding predictions” suffered from the aforementioned issues. This included studies using the L243 or L227 monoclonal antibodies, both anti-DRA, with EBV transformed homozygous (23–27) or heterozygous (28–30) cell lines. Other studies (31, 32) similarly purified DR molecules from heterozygous patient material, yielding a mixture of DRB1 and DRB3 or DRB4 eluted peptides, as well as other undefined DR molecules expressed because of patient heterozygosity.

While most authors do not mention issues pertaining to co-expression, others acknowledge the problem. Koning et al. (33) state that “affinity purification … resulted in the isolation of a mixture of the HLA-DRB1 and the HLA-DRB4-locus product DR53.” Similarly, Hill et al. (34) recognize that DRB1 and DRB4 molecules are tightly linked, but conclude that DRB4 is likely to represent a minor fraction of DR molecules, and is therefore “unlikely to influence results.”

### Most non-predicted ligands are weak binders and/or predicted to bind other HLA molecules

Based on the results above, we addressed whether the 207 non-predicted DRB1*04:01, DRB1*03:01, DRB1*04:02 and DRB1*13:02 ligands could have been eluted from associated and co-expressed DRB3/4/5 molecules, or other contaminant HLA class II. We found 22% (46/207) of the non-predicted DRB1-reported eluted ligands were predicted to bind representative DRB3/4/5 molecules (i.e., DRB3*01:01, DRB3*02:02, DRB4*01:01 and DRB5*01:01).

Next, in light of the observation that several papers had isolated ligands from heterozygous DR samples, we examined whether the remaining 161 non-predicted DRB1 ligands may be predicted to bind other common DRB1 or DQ alleles (16). We found that 81% (130/161) of the non-predicted DRB1-reported eluted ligands were, in fact, predicted to bind other common DR or DQ alleles, suggesting that a wrong HLA might have been assigned to these ligands.

Following an alternative line of investigation, we examined whether some of the apparent discrepancies might be explained by eluted ligands being predicted to have “borderline” affinities, perhaps narrowly missing the 1,000 nM cut-off. Out of a total of 628 non-predicted ligands reported to be eluted from DR molecules, 249 (39.7%) had predicted affinities in the 1,000-2,000nM range **(Supplemental Table S1)**. By contrast, in the case of DQ molecules, of 4927 non-predicted ligands, only 1,372 (27.9 %) were borderline predicted non-binders (i.e., IC50 nM 1,000-2,000nM range).

Taken together, these data suggest that contaminating molecules and/or borderline “near miss” predictions explain a major fraction of discrepancies in the case of DR eluted ligands. However, these explanations were less helpful in the case of DQ eluted ligands.

### Addressing discrepancies between elution data and binding predictions by experimental affinity determinations

Next, we chose representative HLA class II alleles from the set described above, considering the number of reported eluted ligands, frequency of the allele in the general human population, and whether an in-house quantitative binding assay (35) was available. Specifically, we selected HLA-DQA1*05:01/DQB1*02:01, DRB1*04:01 and DRB1*15:01 for further study, all alleles for which less than 75% of ligands were predicted to bind at the 1,000nM level. As a control, we selected DRB1*01:01, an allele for which 95% of ligands were predicted to bind with affinities of 1,000nM or better.

For these four selected alleles, 3,442 eluted peptides with lengths of 15-20 amino acids were available, of which 1,410 (41%) were not predicted to bind at the 1,000 nM level **(Supplemental Table S2)**. We excluded sequences with >80% sequence similarity to reduce redundancy, resulting in a set of 728 peptides. From this set, we selected 30 peptides each for DQA1*05:01/DQB1*02:01 and DRB1*04:01 and 15 peptides each for DRB1*01:01 and DRB1*15:01 (**Table 2**). The number of eluted ligands that were not predicted to bind was smaller for DRB1*01:01 and DRB1*15:01, which is why 15 peptides were selected for testing for these alleles (**Supplemental Table S2**). This resulted in a total of 90 eluted peptide sequences that were not predicted to bind at the 1,000nM level. As a control, we selected 20 eluted peptides per allele that were predicted to bind with affinities of 250 nM or better. Binding affinities for the 90 peptides predicted not to bind and 80 predicted binders were measured using standard competitive binding assays (35), as described in the Methods section. **Supplemental Table S3** lists all tested peptides together with predicted and measured IC50 values.

**Table 2.** Composition of peptide set and associated binding results

| **HLA allele** | **Pred. non-**<br>**binders**<br>**selected** | **Pred. non-**<br>**binders that**<br>**don't bind** | **Pred.**<br>**binders**<br>**selected** | **Pred. binders**<br>**that bound** |
| --- | --- | --- | --- | --- |
| DQA1*05:01/DQB1*02:01 | 30 | 20 (67%) | 20 | 19 (95%) |
| DRB1*01:01 | 15 | 12 (80%) | 20 | 18 (90%) |
| DRB1*04:01 | 30 | 23 (77%) | 20 | 19 (95%) |
| DRB1*15:01 | 15 | 12 (80%) | 20 | 20 (100%) |
| Total | 90 | 67 (74%) | 80 | 76 (95%) |

Overall (**Table 2)**, 76 of the 80 predicted binders (95%) bound the corresponding HLA molecule when tested experimentally (shown as squares in **Figure 1**), consistent with previous data (22), and as summarized above. Similarly, 67 of the 90 eluted peptides (74%) predicted not to bind (**Table 2)** indeed did not bind when tested experimentally (shown as circles in **Figure 1**). Thus, these results confirmed the general efficacy of the binding predictions, as applied to the experimentally determined binding capacity of reported eluted ligands.

**Figure 1.**
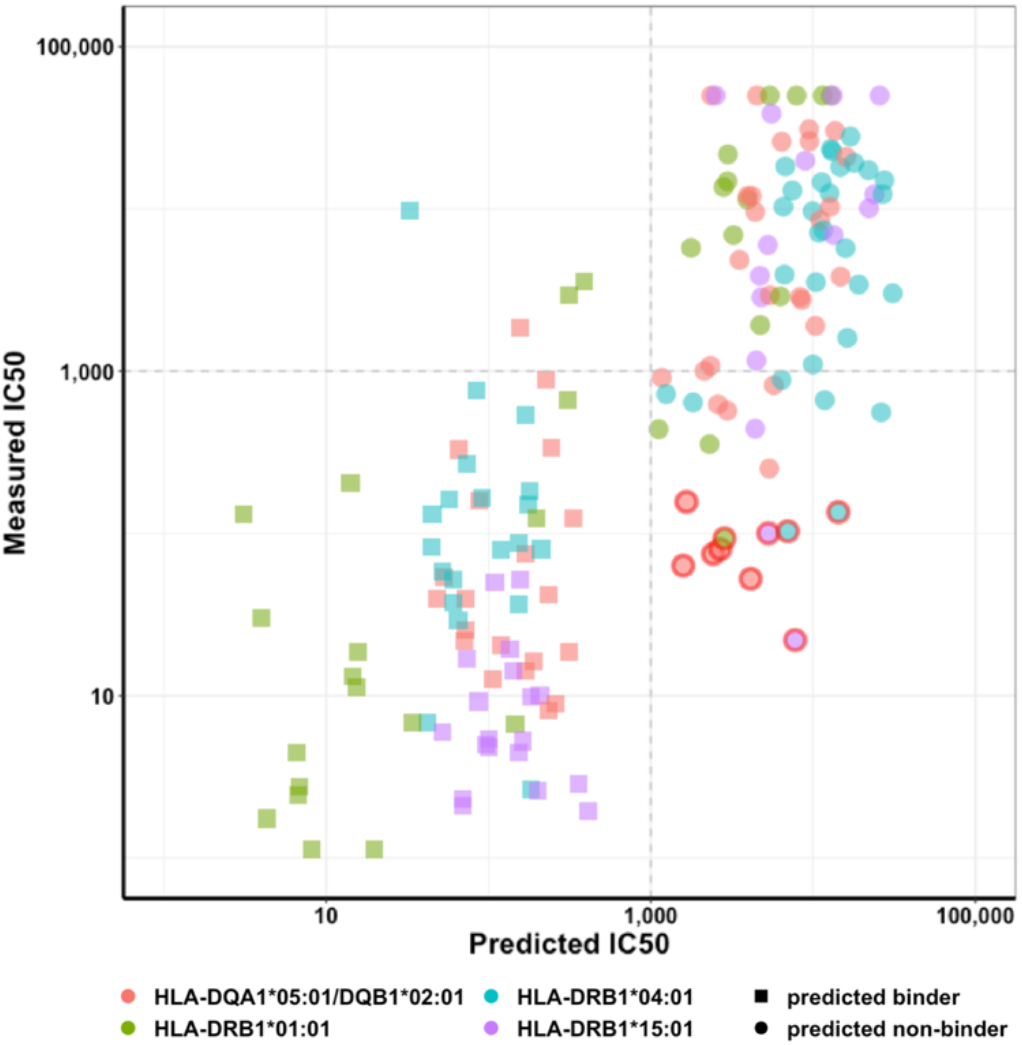
Measured IC50 values are plotted against predicted ones. Colors indicate different alleles, squares predicted binders, and circles predicted non-binders. A red outline depicts cases with large discrepancies (> 10-fold) between measured and predicted affinities.

Importantly, for some alleles a relatively large fraction of eluted peptides does not, in fact, bind the reported corresponding allele when measured experimentally. In light of the complexity and limitations of eluted ligand determinations, it appears likely that at least some of these reported ligands have an incorrectly assigned HLA molecule, or are otherwise contaminants.

### Correlation between predicted and measured binding is lowest for DQA1*05:01/DQB1*02:01

Comparison of the NetMHCIIpan predicted IC50 values to the measured affinities shown in **Figure 1** reveals overall good correspondence. As noted above, most (74%) peptides that were predicted to not bind did in fact not bind when tested experimentally (shown as circles in **Figure 1**), and 95% of the predicted binders did bind, as expected. However, this analysis also identified several cases of disagreement between predicted and measured binding. In particular, the analysis identified 9 peptides (outlined in red in **Figure 1**) that were not predicted to bind but had a 10-fold or higher measured binding affinity than predicted. This suggests that these peptides are likely real ligands whose binding capacity was missed by the predictions. Further inspection of the affected peptide-HLA pairs showed that the majority of these peptides (5 of 9) were associated with DQA1*05:01/DQB1*02:01 (**Table 3**).

**Table 3.** Peptides showing greatest discrepancy between predicted and measured affinity

| **Protein** | **Pos** | **Sequence** | **HLA allele** | **Measured**<br>**IC50** | **Predicted**<br>**IC50** | **Fold-**<br>**change** |
| --- | --- | --- | --- | --- | --- | --- |
| SR | 812 | RQEEPEYENVVPISRPP | DQA1*05:01/DQB1*02:01 | 52 | 2420 | 47 |
| LAM | 246 | KPPTADLFTGVLPNGYNPP | DQA1*05:01/DQB1*02:01 | 117 | 2681 | 23 |
| Cochlin | 95 | LPGRENYSSVDANGIQ | DQA1*05:01/DQB1*02:01 | 445 | 4121 | 9.3 |
| LAT2 | 21 | GEPDYVNGEVAATEA | DQA1*05:01/DQB1*02:01 | 38 | 1582 | 42 |
| Tubulin | 38 | GDSDLQLDRISVYYNEA | DQA1*05:01/DQB1*02:01 | 162 | 1659 | 10 |
| X orf 26 | 23 | SLPAESYGNDPDIEMAWAM | DRB1*01:01 | 93 | 2847 | 31 |
| NFkB | 238 | SRLPGPTGSVVSTGTSF | DRB1*04:01 | 103 | 6986 | 68 |
| 60s rp | 58 | STAAAPAEKKVEAK | DRB1*04:01 | 135 | 14303 | 106 |
| B actin | 356 | WISKQEYDESGPSIVHRK | DRB1*15:01 | 100 | 5301 | 53 |

Accordingly, we next sought to characterize the respective modes of binding of the 5 ‘discordant’ peptides to DQA1*05:01/DQB1*02:01 (hereafter DQ2.5). As controls, we also included 2 peptides eluted from DQ2.5 that were predicted to bind. We also included one peptide (aTIP 430) previously reported as a DQ2.5 restricted T cell epitope (36) and predicted to bind DQ2.5, and another (DQA 16) previously identified as an endogenously bound DQ2.5 ligand (37), but not predicted to bind DQ2.5. The DQ2.5 binding capacity of both aTIP 430 and DQA 16 was previously characterized using purified HLA (17, 38), and confirmed in the present study. **Table 4** shows the 9 peptides included in the single amino acid substitution experiments. **Table 3** above lists the 9 peptides with greatest discrepancy between predicted and measured affinity, where an enrichment for DQ2.5 was noted (5/9). Some peptides are listed in both Tables, as they are relevant in both contexts.

**Table 4.**
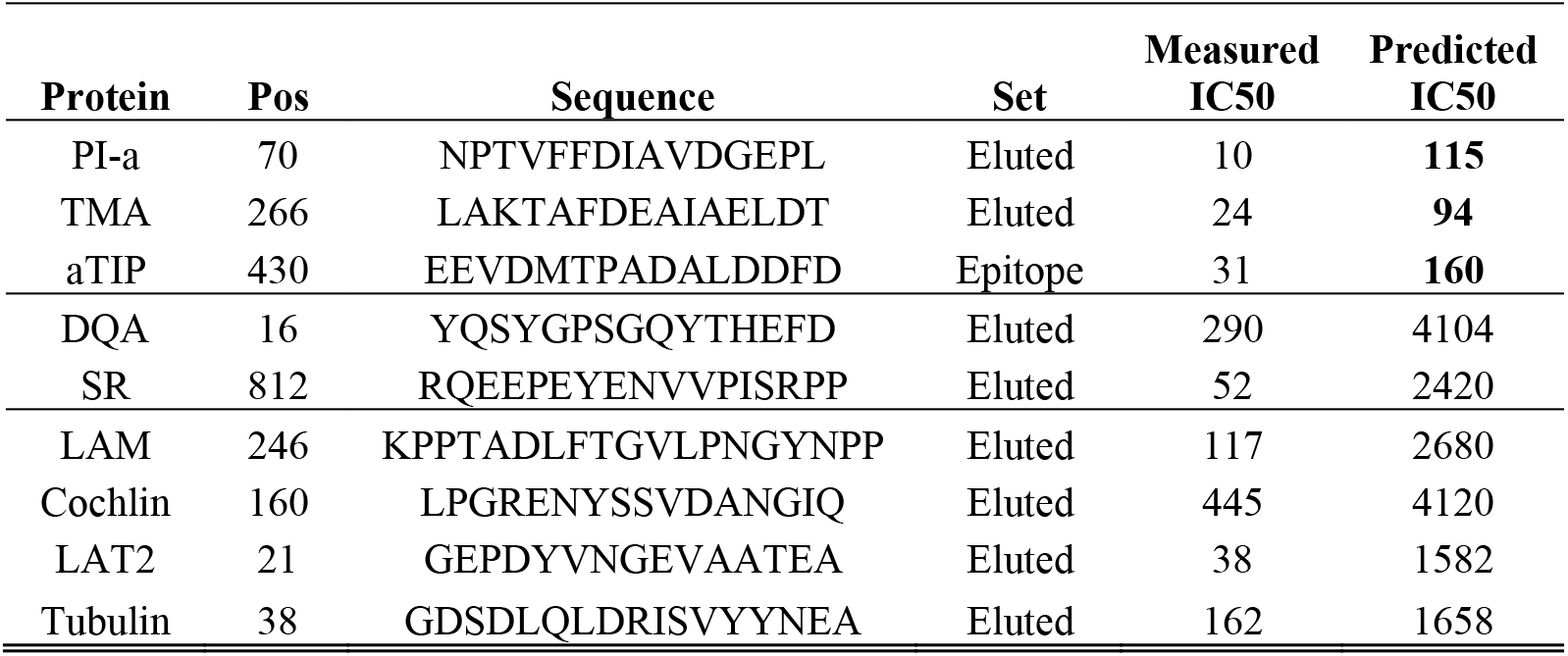
Peptides selected for DQ2.5 single amino acid substitution scan to determine the residues crucial for DQ2.5 binding

### Determining key residues for DQ2.5 binding within the eluted ligands

For each of the ligands in **Table 4** we experimentally determined the residues crucial for conferring DQ2.5 binding capacity. We used a previously described single amino acid substitution approach (39), scanning each peptide sequence by introducing non-conservative substitutions at each position. Lysine (K) was selected as the basis for the analysis as it is non-conservative with all amino acids except those with a positive charge (i.e., R, H and K). In cases where the native residue carried a positive charge, a glutamic acid (E) substitution was utilized. Each single substitution analog, and the original unmodified peptide, was tested for its capacity to bind DQ2.5. All peptides and the measured IC50 values are detailed in **Supplemental Table S4.**

All of the original unmodified peptides bound with IC50 values <1,000 nM. However, several of the single substitution analogs did not bind at all, or bound with a considerably lower IC50 when compared to the corresponding unmodified peptide. The measured binding affinity of each analog was compared to the unmodified peptide, and a relative binding capacity was calculated as the fold change in IC50 between the peptide and its analog (**Figure 2**). Positions associated with a 4-fold or greater reduction in binding capacity (highlighted by green shading in **Figure 2**) were defined as potential MHC contacts. These decreases in affinity allow mapping potential MHC contact residues for each ligand. The analysis revealed that on average 5.8 residues/ligand were identified as important for DQ2.5 binding, with a range from 2 to 9 residues. No clear, simple, common pattern was readily apparent across ligands.

**Figure 2.**
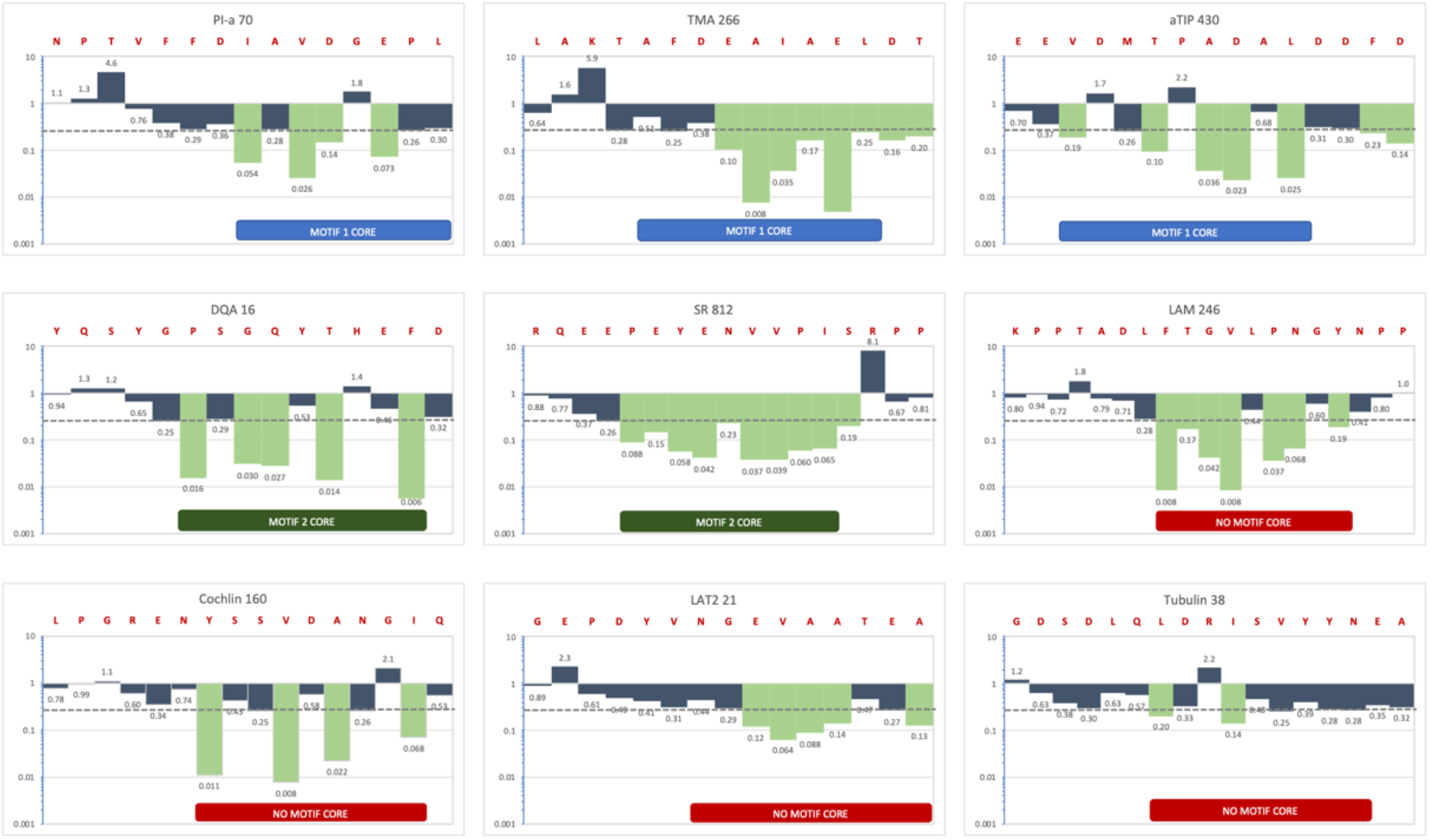
The fold change in IC50 between the unmodified peptide and the corresponding analog with substitutions at each position scanning the length of the respective peptide is shown. The dashed line indicated a 10-fold reduction in binding. Residues for which substitutions resulted in a greater than 10-fold reduction in binding were defined as anchor residues and are highlighted in green here.

### DQA1*05:01/DQB1*02:01 Motif 1, but not Motif 2, peptides are effectively predicted

Previous studies indicated that DQ2.5 may be associated with at least two different binding motifs. One motif is characterized by a preference for aromatic or hydrophobic aliphatic residues in position 1 (P1) (17, 37, 40), acidic residues in positions 4, 6 and/or 7 (P4, P6 and/or P7), and hydrophobic or aromatic residues at position 9 (P9). An independent study (37) also identified a preference for acidic residues in P4 and P6, and large hydrophobic residues in P9, but with no specificity in P1. Collectively, we hereby refer to these preferences as “Motif 1”. Other reported patterns (41, 42) associate proline or polar residues with P1, acidic or polar residues at various other positions, and a hydrophobic or polar residue in P9 (“Motif 2”).

Upon closer inspection of the data shown in **Figure 2**, we were able to classify five ligands as fitting either Motif 1 or Motif 2 (**Figure 3**). Specifically, three peptides appear to fit Motif 1, to include PI-a 70 with isoleucine at peptide position 8 (I8) as its P1 anchor, and aspartic acid at position 11 (D11) and glutamic acid at position 13 (E13) as P4 and P7 anchors, respectively. Motif 1 also describes the binding of the TMA 266 ligand, with glutamic acid at peptide position 8 (E8) and leucine at position 13 (L13) as the P4 and P9 anchors, and aTIP 430, with valine at position 3 (V3), aspartic acid at position 9 (D9), and leucine at position 11 (L11) as P1, P7 and P9 anchors, respectively.

**Figure 3.**
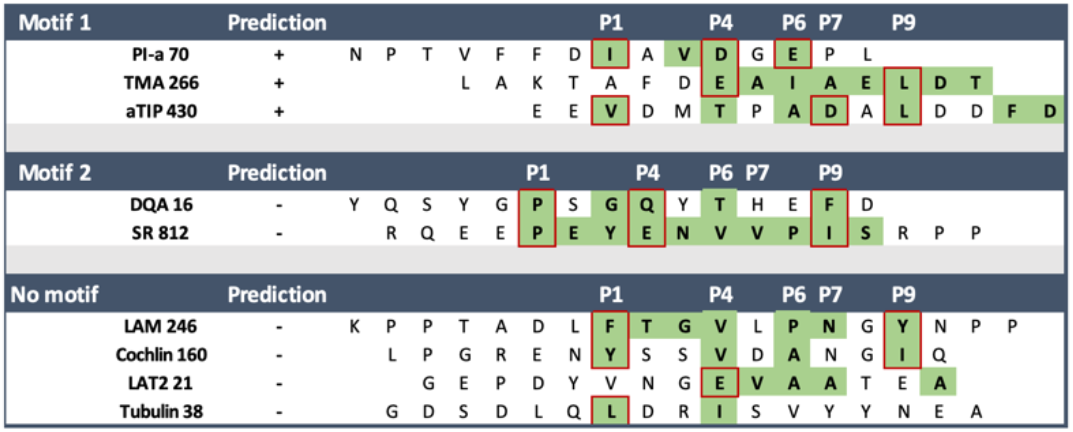
The analyzed 9 DQ2.5 binding peptides are grouped according to the motif they conform to. Residues that were identified as anchor residues in single amino acid substitution experiments are highlighted in green. The red boxes highlight residues that explicitly match the position and specificity of Motif 1 or Motif 2. A + in the column prediction indicates that the peptide was predicted to bind DQ2.5 with an IC50 of < 1,000nM.

Two other peptides appear to fit Motif 2 (**Figure 3**). These are the DQA 16 peptide, for which the single substitution analysis indicated the proline in position 6 of the peptide (P6) as the P1 anchor and phenylalanine at position 14 (F14) as the P9 anchor, and the SR 812 ligand where the proline in position 5 (P5) is the P1 anchor and isoleucine at position 13 (I13) is the P9 anchor.

The four remaining peptides did not fit either motif. These include the LAM 246 and Cochlin 160 peptides, which have aromatic/hydrophobic residues that fit as P1 and P9 anchors, but do not have acidic residues in the P4, P6 or P7 positions as described by Motif 1; meanwhile, aromatic residues in P1 render the LAM 246 and Cochlin 160 peptides incompatible with Motif 2. For the remaining two peptides (LAT2 21 and Tubulin 38) the single amino acid substitution scan also revealed binding modes not compatible with either Motif 1 or 2.

Interestingly, when we examined the motif groups with respect to predicted binding, it was found that all three peptides fitting Motif 1 were also predicted by the NetMHCIIpan algorithm to bind DQ2.5 with an affinity of 1,000 nM or better (**Table 4**). Conversely, none of the peptides associated with Motif 2 or with no motif were predicted by the NetMHCIIpan algorithm. In conclusion, while 3/3 Motif 1-positive peptides were predicted, 0/6 peptides that did not carry this motif were predicted (p=0.019, Fisher’s exact test).

To validate that the differential predictability of DQ2.5 ligands was associated with the presence of Motif 1, we assembled all described DQ2.5 eluted ligands and evaluated whether those predicted to bind DQ2.5 were enriched for the presence of Motif 1, compared to ligands that were not predicted to bind. Indeed, 54% (502/928) of the ligands predicted to bind did carry Motif 1. By contrast, only 26% (298/1127) of ligands not predicted to bind carried Motif 2 (p<0.0001, Chi-square test). The remaining 20% of the ligands predicted to bind DQ2.5 did not conform to any known motif. The neural network architecture utilized by NetMHCIIpan theoretically has the capacity to learn and predict multiple binding motifs for an allele. Our results, however, suggest that the algorithm in practice predicts Motif 1 more efficiently.

Taken together, these data highlight that DQ2.5 can bind peptides promiscuously using alternate modes. Further analysis of more peptides is necessary to fully understand what defines the way peptides bind or do not bind this molecule and to improve binding predictions. Currently, peptides utilizing Motif 1, characterized by the presence of hydrophobic residues in position 1 and/or 9, and acidic residues in positions 4, 6 and/or 7, are being predicted more efficiently.

## DISCUSSION

In this study, we assessed how well MHC class II binding predictions and ligand elution data agree with each other. We found that the majority of experimentally identified MHC class II eluted ligands were predicted to bind to their reported restriction element. For HLA-DR ligands that showed discrepancies, most of the ligands that were not predicted to bind were actually predicted to bind just below the threshold utilized, or were predicted to bind to other co-expressed MHC class II molecules.

In contrast to the DR molecules studied, we found that for HLA-DQA1*05:01/DQB1*02:01 (DQ 2.5) binding predictions did miss several peptides that were determined experimentally to be binders. By determining key residues for DQ2.5 binding in several peptides, we could trace discrepancies back to the existence of two binding motifs, only one of which was effectively predicted.

Our finding that the majority of eluted ligands were predicted to bind their restricting allele supports the use of these prediction tools that are commonly used (43, 44). This use had been put into doubt by reports that the predicted binding affinity of peptides to MHC class II correlates poorly with MHC antigen presentation (12, 13) and peptide immunogenicity (14, 15). We have reported previously that in the case of HLA class II restricted T cell epitopes binding predictions do perform well, as we found that 83.3% of epitopes are predicted to bind their corresponding restricting allele with an affinity of 1,000 nM or better (22).

In the present study, we investigated how efficiently binding predictions can predict eluted ligands, and found that, on average, 64% of eluted peptides are predicted to bind their corresponding allele at the 1,000nM level. This data proves that the majority of eluted ligands can also be effectively predicted, although the percentage is lower than it is in the case for HLA class II restricted T cell epitopes.

Our finding that several HLA-DR alleles presented ligands with predicted affinities in the 1,000-2,000nM range indicated that the 1,000nM threshold might not be accurate for all HLA class II alleles. It was already shown in the case of HLA class I restricted T cell epitopes that alleles can vary in terms of the affinity threshold associated with immunogenicity (45). This may indeed be the case as well for HLA class II ligands and T cell epitopes, and needs to be further investigated.

Our study utilizes experimentally eluted MHC ligand data as a set of ‘positive data points’, and examines which of these ligands is not identified with prediction methods, thus focusing on ‘false negative predictions’. Examining the flip-side, namely ‘false positive predictions’ requires ligand elution datasets that exhaustively search for all possible MHC ligands in a set of antigens so that a set of ‘negative data points’ that with certainty are not presented by MHC can be identified. The datasets used in this study do not meet these criteria, and generating them in sufficiently large scale is challenging. For MHC class I, a comprehensive dataset of eluted ligands derived from vaccinia virus infected murine cells that were further tested for T cell recognition has been published, revealing 83 major epitopes (46). Benchmarking revealed that the best prediction methods were able to capture more than 50% of these epitopes in the top N = 277 predictions within the set of N = 767,788 possible peptides(47). This means the false positive rate, defined as the fraction of negatives that is incorrectly predicted to be a positive is only 0.03%. While this seems close to perfect, when evaluating this performance based on precision, defined as the fraction of positive predictions made that is actually positive, is only 15%. This highlights that metrics of prediction performance have to be considered in the context of the problem considered, and identifying MHC ligands and/or T cell epitopes in a complex antigen is a ‘needle-in-the-haystack’ problem. It also has to be pointed out that the prediction methods were asked to identify ligands based on the peptide sequence alone, and other factors that can lead to lack of presentation of a ligand, such as low expression level of the source protein, were not included in this comparison. Incorporating such additional factors should reduce the false positive rate. Similar datasets and benchmark comparisons should also to be generated for MHC class II.

When we scrutinized experimental procedures of the ligand elution studies we found that the assigned restrictions for several MHC class II ligands are likely incorrect. This reflect. experimental challenges of ligand elution assays. The first steps in a typical MHC class II ligand elution assay are lysing the antigen presenting cells, purifying the MHC molecules via immunoprecipitation, and chromatographically eluting the MHC ligands (48). Here, the main drawback is that natural ligandomes are multi-allelic. Different MHC molecules are expressed in the interrogated cell and the eluted ligands can correspond to any one of those MHC molecules. This makes it difficult to unambiguously annotate the ligands to the MHC molecule they were presumed to be eluted from.

Indeed, we have shown here that in the case of several HLA-DR studies, it was ignored that DRB1 and DRB3/4/5 are co-expressed, and that the HLA annotation of the reported ligands was thereby prone to some degree of ambiguity. This issue can be addressed, for example, by using genetically engineered mono-allelic cell lines. This approach, however, is not feasible when patient samples are used, which will almost always be multi-allelic. To overcome this, there are currently significant efforts to develop computational methods that can accurately annotate multi-allelic ligands to their respective MHC (49, 50). However, such methods will still rely on the identification of shared motifs of the eluted peptides.

Once ligands are eluted from MHC molecules, they are sequenced by tandem mass spectrometry (MS/MS). A key step is the identification of peptides by matching acquired MS/MS spectra against a customized protein sequence database. This step has an inherent trade-off between sensitivity and specificity, which leads to a certain number of false spectrum identifications (51, 52). It is thus expected that some ligands identified by these methodologies are not correct. Given that we explicitly focused on peptides that were not predicted to bind, we might have enriched for such peptides that fall into the known false discovery rate of peptide elution experiments.

On the other hand, specifically focusing on these outliers might have enabled us to discover patterns that would have otherwise been missed. When we compared predicted and measured affinities we identified several peptides that were not predicted to bind their associated allele but were experimentally determined to actually bind. This, therefore, indicates that these peptides are likely real ligands, but were missed by the predictions. As 5 out of 9 of these instances were associated with HLA-DQA1*05:01/DQB1*02:01 (DQ2.5), we performed a detailed investigation of binding modes to DQ2.5. We performed single amino acid substitutions for these 5 discordant DQ2.5 associated peptides and an additional 4 peptides that are known DQ2.5 binders. Two distinct binding motifs have been described for this allele in the literature, and 5 out of the 9 analyzed peptides were determined to fit either of these two motifs, while 4 peptides did not fit any motif.

It might be that these peptides that did not fit any known motif bind DQ2.5 in a mode engaging pockets in the binding groove in a not readily apparent alternate mode. But, as suggested by several elegant thermodynamic studies (53–56), it is also possible that the primary energy of binding in these cases is driven by MHC backbone-peptide side chain interactions. However, without additional data, this remains speculation in the present case.

Interestingly, only peptides conforming to Motif 1 could be predicted correctly as binders with affinity <1,000nM. Peptides conforming to Motif 2 or no motif were not predicted correctly, indicating that prediction performance might depend on the motif of the peptide in question.

There is already an example in the literature of an allele having two alternative motifs (57). Interestingly, this is HLA-DRB1*03:01, which was also found in our study to be associated with lower prediction performance, with only 52% of peptides binding at the 1,000nM level.

It is possible that alternative binding motifs are a general problem with HLA class II molecules and that currently only peptides conforming to one of the associated motifs are predicted well. Conversely, peptides with other motifs are missed by predictions. While the neural network architecture utilized here hypothetically has the capacity to learn that multiple binding motifs are possible for a single allele, the algorithm in practice does not seem to pick this up. This needs to be investigated in more detail by extending the analysis to additional alleles and peptides. The resulting dataset from our study will be valuable for training machine learning methods to specifically detect cases with multiple binding motifs and improve prediction performance.

DQ2.5 has been specifically identified as a primary genetic risk factor for celiac disease(58). Binding of deamidated gliadin peptides, which are components of gluten, leads to activation of gluten-sensitized T cells, which triggers the autoimmune response. The deamidation by tissue transglutaminase converts glutamine residues to glutamic acid, which makes the gliadin peptide negatively charged(59). This negative charge facilitates the binding to HLA-DQ2.5, as the P4, P6, and P7 pockets have a preference for negatively charged residues (37, 60). Motif 1 and Motif 2 share a preference for negatively charged residues;. As gliadin peptides are proline rich, they more generally conform to Motif 2, which has a preference for proline as the P1 anchor residue.

Crystallography studies of DQ2.5 with bound peptides conforming to Motif 1 and Motif 2 would enable the detailed characterization of the interactions in the two binding modes. Nine crystal structures of DQ2.5 are available in the Protein Database (PDB). However, as DQ2.5 is mainly studied in the context of celiac disease, in 7 of these studies various deamidated gliadin peptides are reported, which all conform to Motif 2 (61–63). The remaining 2 crystal structures report CLIP1 and CLIP2 peptides, which do not conform to any motif (64).

Crystal structures of DQ2.5 with bound gliadin peptides or mimics of these peptides in complex with T cell receptors (TCRs) showed that the TCR-peptide contacts are at positions P2, P5, P6 and P8 of the MHC binding core (62, 65). Both Motif 1 and Motif 2 do not identify a preference for specific amino acids at positions P2, P5, and P8, and have the same preference for acidic residues at positions P6 and P7. It is hence theoretically possible that peptides conforming to Motif 1 or Motif 2 could be recognized as epitopes. Again, crystal structures of DQ2.5 with bound peptides conforming to Motif 1, ideally in complex with a TCR, would help to clarify the specific interactions of the two binding motifs.

## MATERIAL AND METHODS

### Data assembly

The Immune Epitope Database (IEDB)(66) catalogs experimental data on antibody and T cell epitopes as well as MHC ligands. The IEDB was queried to retrieve ligand elution data for MHC class II using the following criteria: “Positive Assays Only, Epitope Structure: Linear Sequence, No T cell assays, No B cell assays, MHC ligand assays: MHC ligand elution assay, MHC Restriction Type: Class II.” The collected data included ligand sequence, the MHC class II allele from which the ligand was eluted from, details of the source protein from which the ligands originated from, and PubMed identifiers (PMIDs) of the studies that reported the ligand.

We further refined this set to only include peptides that were eluted from HLA molecules typed with 4-digit resolution, resulting in a set of 24,601 peptides from 27 different HLA class II molecules. The length of the ligands in this set ranged from 3 to 43 residues; based on previous analyses (22) we only retained peptides of 15-20 amino acids in length. After this step, the dataset contained 12,506 peptides from 27 different HLA class II molecules. We then further filtered the set by considering only alleles for which at least 10 ligands were reported. The resulting final dataset used for analysis included 12,449 sequences, eluted from 16 different HLA class II molecules. The composition of this dataset with peptide numbers per allele are described in detail in **Supplemental Table S1**.

### HLA binding predictions

We performed binding predictions using the standalone version of NetMHCIIpan (version 3.1), as this tool provides predictions for most HLA class II alleles (20, 21). HLA class II binding predictions were performed for all 12,449 sequences and their reported HLA class II restriction element. Peptides with predicted binding affinities of >1,000nM were considered predicted non-binders. This threshold was selected based on previous studies analyzing the binding affinity of HLA class II restricted T cell epitopes (16–19).

### Selection of a peptide set with discrepancies

We selected 4 alleles to further investigate peptides with discrepancies between ligand elution data and binding predictions: HLA-DQA1*05:01/DQB1*02:01, DRB1*04:01, DRB1*15:01, and DRB1*01:01. These representative alleles were selected considering the number of reported eluted ligands, frequency of the allele in the general human population, and whether an in-house quantitative binding assay (35) was available. In total, 3,442 eluted peptides were available for the 4 selected HLA class II alleles, of which 1,410 had predicted affinities >1,000nM. Accordingly, these were considered as ‘predicted non-binder’. The composition of the dataset after each filtering step is described in detail in **Supplemental Table S2**.

### Analysis of sequence similarity

Many of the eluted peptides are very similar to each other as they contain the same MHC class II binding core with varying flanking residues of different lengths. When selecting peptides for experimental testing we wanted to avoid testing the same MHC class II binding core multiple times. Thus, to reduce redundancy, we removed sequences with >80% sequence similarity. We used the IEDB clustering tool (67), which groups the set of input peptides into clusters based on sequence identity. Here, a cluster is defined as a group of sequences that have a similarity greater than the specified minimum sequence identity threshold (80% in our case). We used all 1,410 eluted peptides for the 4 selected alleles as input, and from each cluster we randomly selected one peptide. As a result, 728 peptides that had <80% sequence similarity remained in the dataset, from which peptides were selected for experimental testing.

### HLA binding measurements

Purification of MHC molecules by affinity chromatography was performed as detailed elsewhere(35). Briefly, Epstein-Barr virus (EBV) transformed homozygous cell lines or single MHC allele transfected RM3 or fibroblast lines were utilized as sources of HLA class II MHC molecules. MHC Class II molecules were purified from cell pellet lysates by repeated passage over Protein A Sepharose beads, conjugated with the monoclonal antibodies (mAbs) LB3.1 (anti-HLA-DR), SPV-L3 (anti-HLA-DQ) and B7/21 (anti-HLA-DP). Protein purity, concentration, and the effectiveness of depletion steps were monitored by SDS-PAGE and BCA assay.

Classical competition assays to quantitatively measure peptide binding to MHC class II molecules were performed as detailed elsewhere(35). These assays are based on inhibition of binding of high affinity radiolabeled peptides to purified MHC molecules. Briefly, 0.1-1 nM of radiolabeled peptide was co-incubated at room temperature or 37°C with purified MHC in the presence of a cocktail of protease inhibitors. Following a two- to four-day incubation, MHC bound radioactivity was determined by capturing MHC/peptide complexes on MHC locus specific mAb coated Lumitrac 600 plates (Greiner Bio-one, Frickenhausen, Germany). We then measured bound counts per minute (cpm) using the TopCount (Packard Instrument Co., Meriden, CT) microscintillation counter. The concentration of peptide yielding 50% inhibition of binding of the radiolabeled peptide was calculated. Under the conditions utilized, where the concentration of the radiolabeled peptide is less than the concentration of MHC, and the affinity of the peptide (IC50 nM) is greater than or equal to the concentration of MHC (or more formally, [label]<[MHC] and IC50 ≥ [MHC]) measured IC50 values are reasonable approximations of true equilibrium dissociation constants (Kd) (68, 69). Each competitor peptide was tested at six different concentrations covering a 100,000-fold range, and in three or more independent experiments. As a positive control, the unlabeled version of the radiolabeled probe was also tested in each experiment.

## Abbreviations

MHC: Major Histocompatibility Complex
HLA: Human Leukocyte Antigen
IEDB: Immune Epitope Database
TCR: T Cell Receptor
PDB: Protein Database
DQ2.5: DQA1*05:01/DQB1*02:01
MS: mass spectrometry
P1, P2, P3, P4, P6, P7, P8, P9: binding pockets 1-9 in MHC binding groove
mAb: monoclonal antibody
EBV: Epstein-Barr Virus
nM: nanomolar
Kd: equilibrium dissociation constant

## Footnotes

Research reported in this publication was supported in part by the National Institute of Allergy and Infectious Diseases of the National Institutes of Health under Award Number R21AI134127 and 75N93019C00001, as well as the National Institute of Dental and Craniofacial Research and the National Cancer Institute under Award Number U01DE028227. The content is solely the responsibility of the authors and does not necessarily represent the official views of the National Institutes of Health. The work was also supported in part by a research agreement with Bristol-Myers Squibb Company.

